# Microbial stem cells support productivity in dedicated factory cells in an asymmetrically dividing *E. coli* system

**DOI:** 10.1101/2025.10.01.679865

**Authors:** Nikolai Mushnikov, Jacob T. Guidry, Ashley Park, Grant R Bowman

## Abstract

A major challenge for many bio-manufacturing operations is that cells are burdened by the high fitness cost of product synthesis, limiting their growth and productivity. A potential solution is to decouple cell reproduction from product synthesis by dividing these conflicting tasks between two differentiated cell types. This work describes the use of an asymmetrically inherited protein cue to differentiate an *E. coli* culture into reproductive stem cells and fully dedicated factory cells. Cell differentiation is based on the ability to accumulate two factors: a variant of phage-derived T7 RNA polymerase (T7RNAP) and GP2, a peptide that inhibits native host cell RNA polymerase. Activating these two factors in factory cells inhibited growth and focused them on T7RNAP-driven product synthesis. Preventing their accumulation in stem cells allowed this cell type to grow and divide asymmetrically, generating new factory cells in the process. These differentiating cell cultures generated over eight-fold higher target protein titers compared to factory cell-only controls. Because they include a mechanism for preventing leaky T7RNAP-driven gene expression in stem cells and pre-induction cultures, it was possible to generate strains with multiple plasmid-based copies of a cytotoxic target gene, whereas leaky expression made the same plasmid inviable in conventional protein expression strains. This versatile genetic system could be useful for generating higher product titers, particularly in cases where product synthesis causes cytotoxicity.

## INTRODUCTION

Bio-manufacturing is recognized as a promising strategy for the large-scale production of molecules that are useful to human society. Currently, genetically engineered microorganisms are used in a variety of industrial processes, ranging from therapeutic proteins to consumer chemicals and biofuels. While the theoretical potential of bio-manufacturing is essentially unlimited, high operating costs and low product titers limit its spread and impact.

A fundamental limitation of most bioengineering approaches is that the accumulation of biosynthetic enzymes, intermediate metabolites, and final products instigates metabolic conflicts between the engineered biosynthetic pathway and natural mechanisms that support the viability of the host cell. High levels of target gene expression can limit growth by monopolizing cellular resources and energy ^1, 2^, and accumulating enzyme products, either as pathway intermediates or endpoints, can intoxicate host cells ^3^. Over multiple cell generations, production stresses can induce a higher rate of mutagenesis and genetic drift ^4^, resulting in genetic selection that favors the outgrowth of non-productive mutants^5^. These negative effects have been mitigated for specific products by optimizing biosynthetic pathways^6–9^, adapting host cell metabolism^10–14^, and improving bioprocess design^15–17^. However, these approaches are typically product-specific and therefore not easily applied toward the biomanufacturing of multiple classes of molecules.

Here we describe Microbial Stem Cell Technology (MiST) and explore its potential as a broadly applicable tool for increasing the productivity of bio-manufacturing processes. The underlying principle, which separates MiST from most other genetic engineering tools, is the functionalization of multicellularity in microbial cultures. In MiST cultures, product synthesis and the associated metabolic burdens are sequestered within a differentiated sub-population of “factory cells”, while a different sub-population of “stem cells” serves as a progenitor of new factory cells, effectively decoupling cell growth from productivity in a culture that maintains the ability to generate new factories. Methods for establishing asymmetric cell division and cell differentiation in *E. coli* cell cultures have been described for an earlier implementation of the MiST system ^18, 19^ and by others ^20,21^.

Among these engineered systems, the asymmetrically localized determinant that drives cell differentiation is a functionalized variant of PopZ, a polar organizing protein found in *Caulobacter crescentus*^22^ and other species of *Alphaproteobacteria*^23^. In its native contexts, PopZ is a pole-localized scaffold that organizes polar micro-domains by interacting specifically with a subset of diffusible cytoplasmic factors^22^. In engineered contexts, the key biophysical property of PopZ is that it self-assembles into a bulky macromolecular complex^22,24,25^ that is usually localized to a bacterial cell pole because it is excluded by the nucleoid, which fills the cytoplasm in the middle of the cell^26^. Since unipolar foci of PopZ remain stably localized through cell division and for multiple generations thereafter, they have been useful as asymmetric cues for organizing cells or guiding cell differentiation^18^.

In *E. coli* and other bacterial hosts ^27, 28–30^, the most common approach toward achieving a high level of target gene expression is to employ bacteriophage-derived T7RNA polymerase (T7RNAP). The key features of T7RNAP are its high level of transcriptional activity relative to host RNA polymerase ^27,31^ and its high level of specificity for its cognate *P_T7_* promoter, a relatively short DNA sequence that is sufficient for driving T7RNAP-mediated transcriptional activity ^32^. In *E. coli* strain BL-21 (DE3) and its derivatives, including many bio-manufacturing strains^33,34^, the gene encoding T7RNAP is integrated into the chromosome under a chemically inducible promoter ^35–37^, while target genes are normally encoded on a plasmid vector bearing *P_T7_* promoters ^37,38^. Typically, low concentrations of inducer are sufficient to drive high-level target gene expression, making this system attractive for industrial applications.

However, T7RNAP-mediated expression systems have substantial limitations. High levels of target gene expression are achieved at the expense of high metabolic burden, as up to 90% of the protein-synthesis machinery can be devoted to heterologous gene expression^39–41^, rapidly exhausting culture growth and production potential ^5^. On the other hand, the high sensitivity of the system can result in significant levels of leaky target gene expression in the uninduced state, which can be problematic in cases where the biosynthetic pathway produces toxins that induce counterproductive selection pressures ^39,42,43^. Mitigation strategies focused on modifying the promoter sequence for balanced expression and reduced leakiness^44^ have improved product yields, but these are only partial solutions to the inherent conflict between cell growth and productivity. Another approach is to decouple growth from productivity, and this has been achieved by temporally separating culture expansion from product synthesis ^45^, and by funneling cellular resources toward production by silencing the transcription of host genes ^46^. However, these techniques limit production to the population of cells that is formed during initial expansion of the biomass.

In this work, we demonstrate a method for integrating high-level T7RNAP-mediated gene expression into MiST genetic circuitry. Here, normal cellular mechanisms for maintaining cell health and cell division are limited to stem cells, and through asymmetric cell division and terminal cell differentiation, they form a stem cell population, and are decoupled from mechanisms that fully commit factory cell progeny to product synthesis in a co-existing, terminally differentiated, and fully committed factory cell population. By merging MiST with user-defined T7RNAP output, we provide a widely applicable platform for stem cell-supported bio-manufacturing processes and demonstrate the potential of such systems for increasing bio-manufacturing yields.

## RESULTS

### Design principles for improved MiST

To use T7RNAP for driving product gene expression in MiST, it was necessary to establish a system in which T7RNAP is inactive in stem cells and active in factory cells. Our approach was to control T7RNAP stability, by combining a variant of T7RNAP that tolerates C-terminal fusions ^47^ with an 11 amino acid SsrA protein degradation tag ^48,49^. The coding sequence of degradable T7RNAP-ssrA was placed under a vanillate-inducible promoter on a low-copy plasmid (Figure 1C). In *E. coli*, SsrA tags are recognized by the SspB adapter, which delivers SsrA-tagged proteins to ClpXP protease for degradation ^48,50^. To specify degradation of T7RNAP-ssrA in stem cells but not factory cells, we translationally fused SspB to the MiST stem cell determinant, creating SspB-mCherry-PopZ (SCP). To control when SCP is expressed, we placed its coding sequence under an arabinose-inducible promoter on a medium-copy plasmid (creating pBAD-SCP). A third “output plasmid” enabled T7RNAP-driven gene expression by placing an output gene (e.g. GFP) under a T7 promoter (pT7-GFP). We expected this genetic circuitry, designated MiST^P^ because it is entirely plasmid-based, to exhibit arabinose-activated expression of the SCP stem cell determinant, which would be asymmetrically inherited by stem cells and excluded from factory cells. Due to the asymmetric inheritance of SCP, we expected that stem cells would degrade T7RNAP-ssrA, while factory would accumulate T7RNAP-ssrA and express output genes (Figure 1D).

**Figure 1.**
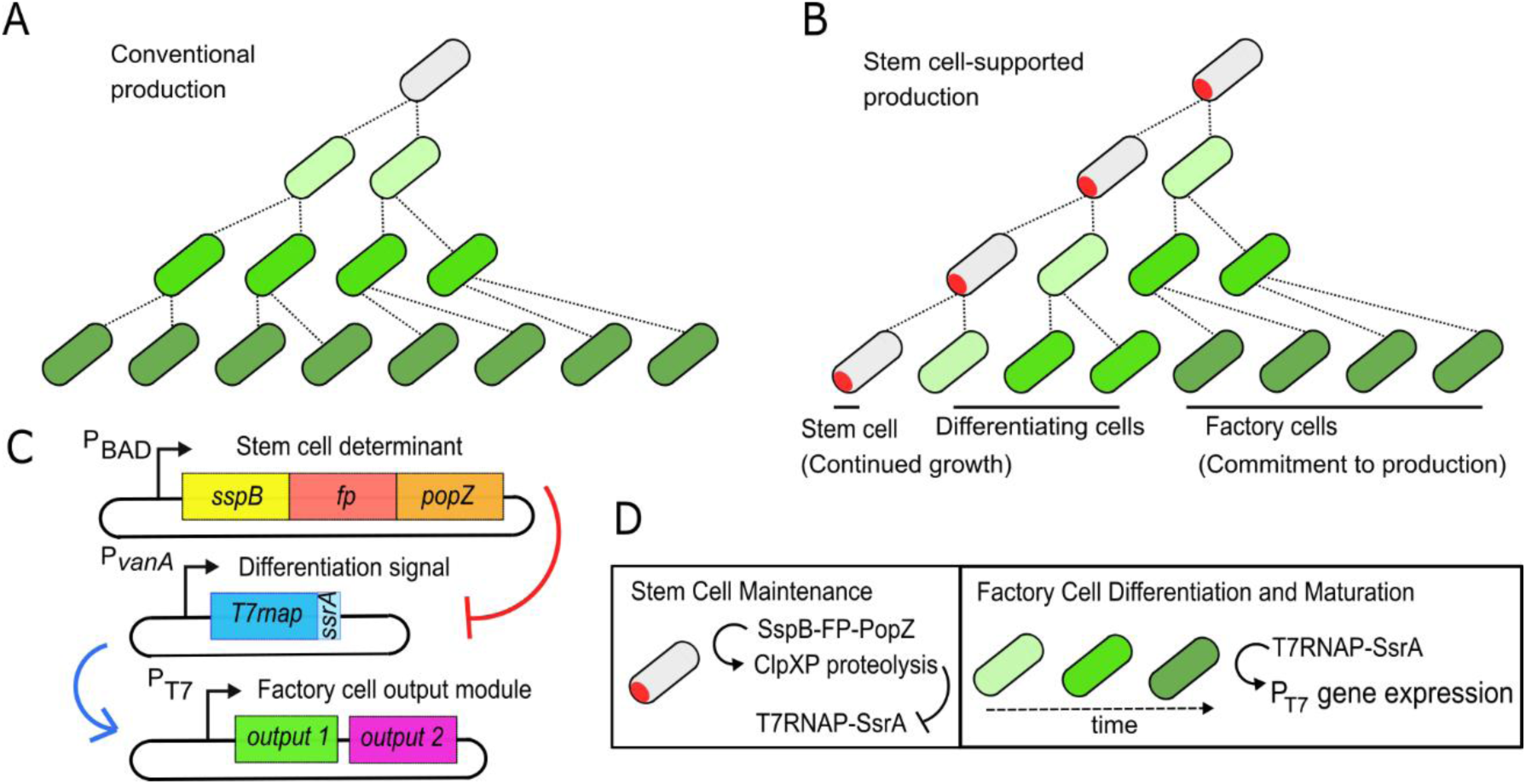
Principles of Microbial Stem Cell Technology (MiST). (A) Conventional bacterial culture: all cells are induced to generate a product (green), which may be coupled to negative effects from biosynthetic burden. (B) Stem cell-supported (MiST) culture: stem cells (red polar foci) divide asymmetrically into one regenerative stem cell and one differentiating factory cell. Factory cells and their progeny generate product (green), and may be affected by biosynthetic burden, while stem cells may be shielded from production stresses. (C) MiST genetic circuitry: (i) Stem cell determinant – PopZ scaffold, functionalized by fusion with SspB adapter for proteolysis, and visualized by fluorescent protein (fp); (ii) Differentiation signal – T7RNAP with ssrA degradation tag on C-terminal end; (iii) Output module – protein(s) of interest encoded under *P_T7_* promoter. (D) Asymmetry and differentiation in MiST cultures. Stem cell identity (left) is maintained by the continual degradation of T7RNAP-ssrA, via SspB adaptor and ClpXP protease activity. Factory cells (right) accumulate T7RNAP-ssrA, which induces target gene expression from *P_T7_* promoter(s).

After demonstrating the functionality of our T7RNAP-ssrA construct (Supplementary Figure 1), we created a complete MiST^P^ genetic circuit by co-transforming pVan-T7RNAP-ssrA, pBAD-SCP, and pT7-GFP into *E. coli* MG1655 (Figure 2A). In a functional MiST strain, a pulse of expression of the stem cell determinant initiates stem cell identity, and following the pulse, the stem cells divide asymmetrically into stem cells and differentiating factory cells. Consistent with these expectations, we observed that SCP formed stable pole-localized foci over multiple cell divisions, and these cells maintained stem cell identity by remaining off for GFP expression.

**Figure 2.**
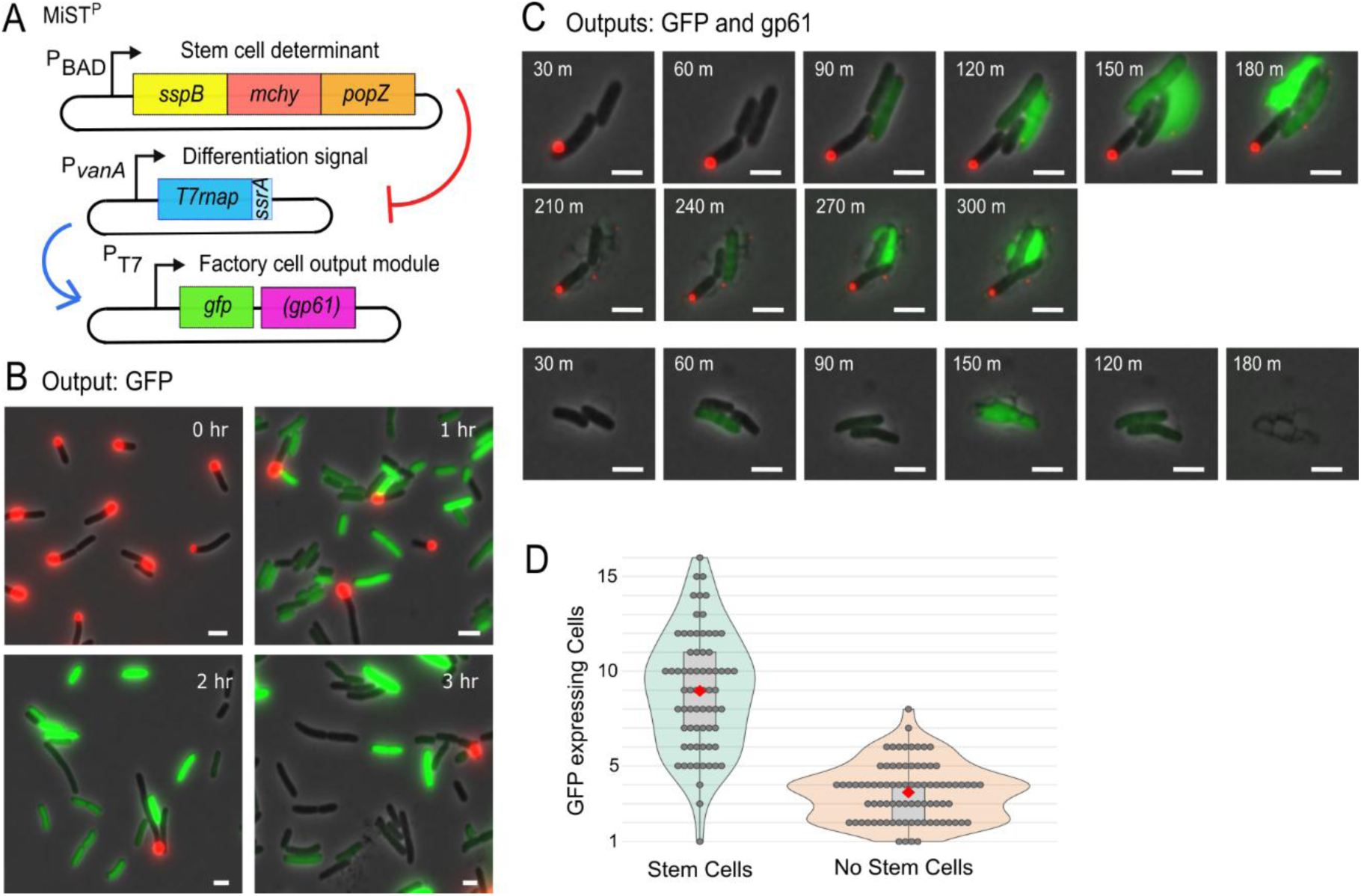
Factory cell differentiation in MiST cultures and protection of stem cells from a toxic product. (A) MiST^P^ genetic circuit, with T7RNAP-ssrA driving expression of GFP and, where indicated, the murein hydrolase gp61. (B) Fluorescence images of MiST^P^ cells after the induction of factory cell differentiation at t = 0 hr. SGP stem cell determinant (red polar foci) and factory cell product (green GFP signal) are overlayed on a phase contrast image (gray). Scale bars 2µM. (C) Time-lapse fluorescence images of stem cell-supported (upper panels) and factory cell-only (lower panels) microcolonies in MiST^P^ cultures, wherein T7RNAP-ssrA drives expression of GFP and gp61. Image series begin shortly after the first cell division following factory cell induction. Cell lysis events are visible in corresponding Supplementary Movie M2. Scale bars 2µM. (D) Total number of GFP-expressing cells generated in stem cell-supported (n=69) and factory cell-only (n=84) microcolonies from C, until complete lysis. Individual data points (gray) are displayed with the population average (red diamonds) and a boxplot showing interquartile range.

Meanwhile, progeny lacking polar foci of SCP differentiated into GFP-producing factory cells (Figure 2B, Supplementary Movie M1). Notably, factory cells exhibited brightly fluorescent output signal within 45 minutes (2 generations) of separation from stem cells, whereas the MiST systems described in our previous work, which rely on the accumulation of c-di-GMP as the factory cell on-switch ^18^, required over 100 minutes for factory cell activation.

### MiST Supports the Production of a Cytotoxic Peptide

To observe the behavior of a MiST cell culture during the production of a cytotoxic molecule, we created a polycistronic output cassette that included the coding sequence of the T4 bacteriophage-derived peptide gp61 downstream of GFP (pT7-GFP-gp61). Gp61 is a murein hydrolase used by the bacteriophage to escape from their bacterial hosts, and its expression quickly results in cell lysis ^51^.

We co-transformed pT7-GFP-gp61 with pVan-T7RNAP-ssrA and pBAD-SCP plasmids to create a MiST^P^ strain. In these cultures, GFP expression was associated with cell lysis (Figure 2C, Supplementary Movie M2). Using time-lapse microscopy to compare the fates of mChy-labeled stem cells to non-stem cells that are free to express T7RNAP-dependent gene products, we observed that non-stem cells acquired bright GFP fluorescence and generated an average of 3.6 descendants before all cells lysed. In contrast, stem cells did not express GFP and generated an average of 8.9 descendants before all cells lysed (Figure 2D). If the number of GFP-expressing descendants is an indicator of factory cell productivity, these results suggest that a stem cell-supported culture would express more GFP and Gp61 than a control culture that expressed the same products without stem cell support.

### Improving the Stem Cell Determinant and Committing Factory Cells to Product Synthesis

We were concerned that MiST^P^ strain performance could be negatively affected by the presence of an alternative translation initiation site near the N-terminus of mCherry’s coding sequence, which could lead to the production of a truncated product ^52^. Consistent with this, western blotting revealed that SCP was expressed in two major forms (Supplementary Figure 2A, 2B). To alleviate this issue, we replaced mChy with GFP, creating an SspB-GFP-PopZ fusion, hereafter referred to as SGP. Western blotting indicated that SGP is expressed as a single fragment, approximately 80kDa in size. To maintain dual-color tracking of Stem and Factory cells, we created a mChy output cassette, pT7-mChy. Inducing the MiST program with these components resulted in expected observations of asymmetric cell division and factory cell differentiation (Supplementary Figure 2C, 2D).

We further improved the system by switching the cytotoxic factory cell output from Gp61 to the T7 bacteriophage-derived peptide Gp2, creating output cassette pT7-mChy-gp2. Since Gp2 is a non-lytic cytotoxin, this alleviated the complication of quantifying yields from lysing factory cells. Moreover, GP2 specifically inhibits bacterial RNA polymerase while allowing T7RNAP to remain active ^53^, which focuses cell resources on T7RNAP-regulated gene expression ^46^. We predicted that the expression of this polycistronic message would arrest cell growth and induce high levels of mChy production (Figure 3A).

**Figure 3.**
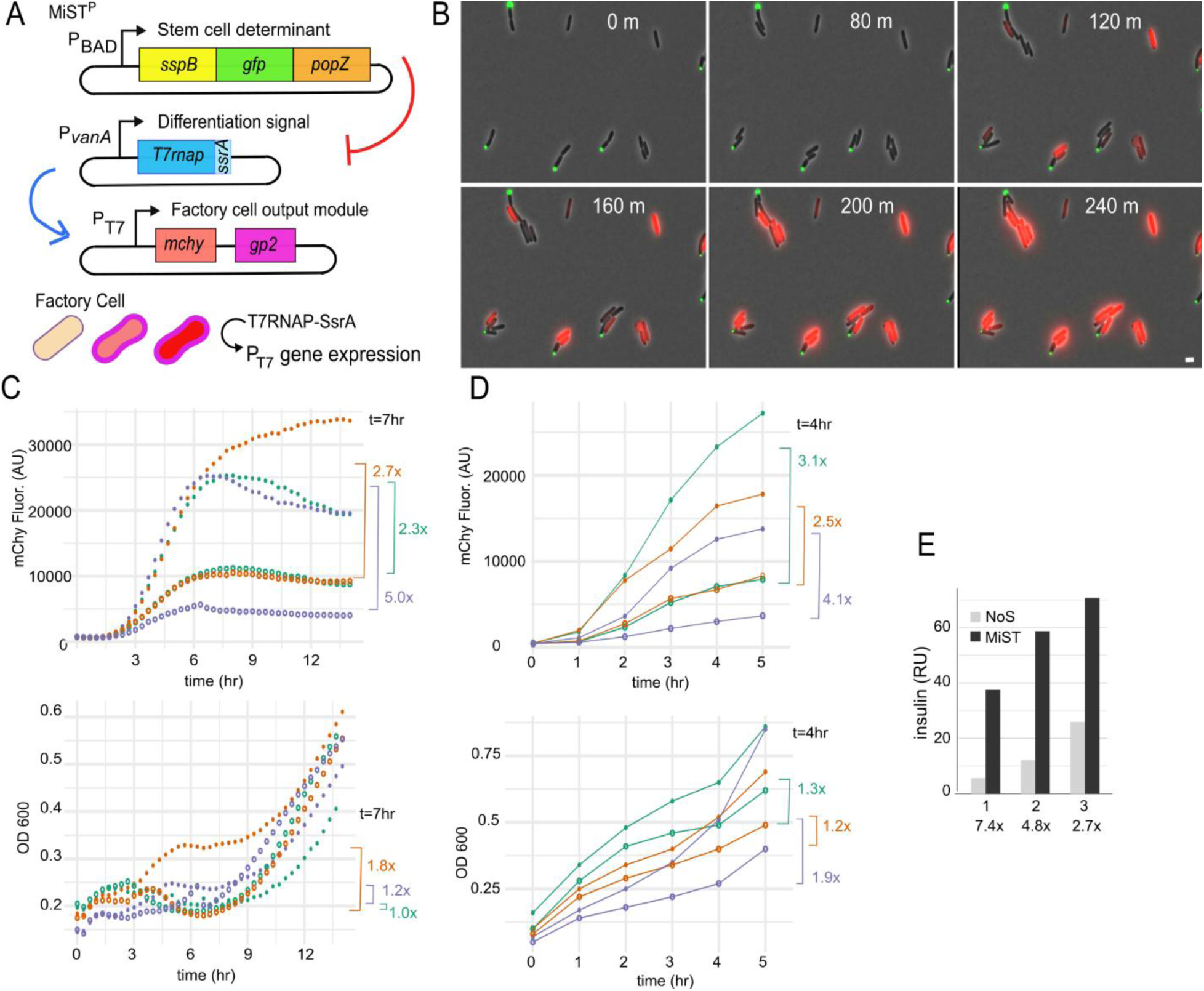
Higher production from dedicated factory cells in stem cell-supported MiST cultures than from factory cell-only cultures. (A) MiST^P^ genetic circuit, with T7RNAP-ssrA driving expression of mCHY and gp2. (B) Time-lapse fluorescence images of MiST^P^ cells after the induction of factory cell differentiation at t = 0 hr. SGP stem cell determinant (green polar foci) and factory cell product (red mCHY signal with gp2, not visible) are overlayed on a phase contrast image (gray). Scale bars 2µM. (C, D) mCHY fluorescence intensity (upper panel) and OD600 (lower panel) in stem cell-supported MiST^P^ cultures (filled circles) versus factory cell-only controls (empty circles), grown in a microplate reader (C) or in 50 ml shake flasks (D). Colors differentiate three separate experiments, with fold differences at the indicated time points denoted by brackets. (E) Insulin titers produced by stem cell-supported (MiST) versus factory cell-only control cultures (NoS) after 20 hours of production, in three independent experiments, with fold differences indicated.

Inducing the MiST program with these components resulted in the growth arrest of factory cells and concomitant accumulation of mChy fluorescence. Meanwhile, stem cells continued to divide and generate new factory cell progeny (Figure 3B; Supplementary Movie 3). We compared stem cell-supported (i.e., MiST) cultures to a control, which was the same strain prepared without stem cell induction, in which all cells were induced to act as factories. When we induced product synthesis in microplates, stem cell-supported MiST^P^ cultures produced, on average, 3.3-fold more mCherry than controls (Figure 3C). To rule out clonal variation as a confounding factor, we re-transformed the plasmid components into fresh strains, creating four independent strains. Again, we observed that stem-cell-supported cultures produced 2.5 to 3.5-fold more mChy than factory cell-only controls (Supplementary Figure 3A).

We measured mChy production and observed cell differentiation in MiST^P^ cultures growing in 50 ml shake flasks. Under these conditions, stem cell-supported cultures exhibited an average of 3.2-fold more mCherry fluorescence signal compared to controls (Figure 3D). We quantified population dynamics in shake flask cultures by plotting the GFP and mChy fluorescence intensities of individual cells (Supplementary Figure 3B). In MiST^P^ cultures, we observed three distinct population groupings: GFP-positive/mChy-negative stem cells, GFP-negative/mChy-positive factory cells, and double-negative cells. The latter category includes newborn factory cells that have not yet had time to differentiate (Supplementary Movie M3). However, “cheater mutants”, which have lost the ability to produce toxic products due to selective pressure against growth inhibition, are also expected to be double-negative for fluorescence. We found that the population fraction of double-negative cells continued to increase even at the 4hr and 5hr time points, when the percentage of stem cells is low, suggesting that cheater mutants have emerged and are proliferating. Consistent with this, the increase in mChy production plateaued at 5hrs, while the optical densities of the cultures began to rise (Figure 3D). The concomitant loss of production and overpopulation of cheater mutants is representative of the challenges of industrial biofermentation ^4^.

To determine whether factory cell enhancement in MiST^P^ cultures could be beneficial in manufacturing a valuable protein product, we added the pro-insulin coding sequence, tagged with a FLAG epitope, into an output cassette, enabling co-expression of pro-insulin with mCherry and Gp2 (pCDF-Ins-mChy-Gp2). Stem cell-supported (MiST) and factory cell-only control cultures were grown in parallel in 50 mL shake flasks, and whole-cell lysates were collected after 20 hours of production. Western blotting with an anti-FLAG antibody showed that MiST^P^ cultures accumulated, on average, 3.2-fold more proinsulin than cultures that lacked stem cell support (Figure 3E).

### Enhanced Productivity in Differentiating Factory Cells

We observed that the average level of mChy accumulation in individual factory cells of MiST^P^ cultures was higher than the average level of mChy in factory cell-only controls (Figure 4A). This suggests that the factory cells that differentiate from stem cell ancestors have a production advantage relative to factories that are derived from undifferentiated precursors. One possible explanation is that the presence of SGP in stem cells primes the newly differentiating factory cell progeny for high levels of expression by promoting rapid T7RNAP-ssrA accumulation. This could occur if SGP forms heterodimeric complexes with native SspB, which is normally a homodimer ^54^, and sequesters it in stem cell polar foci. To investigate SspB localization, we expressed SspB-mScarlett by itself, or with the normal MiST stem cell determinant (SGP), or with a truncated stem cell determinant that is comprised of the first 13 amino acids of SspB fused to GFP-PopZ (xGP) (Figure 4B). When SspB-mScarlett was expressed by itself, it was diffusely distributed throughout the cytoplasm. When co-expressed with xGP, SspB-mScarlett faintly co-localized with the polar marker, suggesting a weak association. In contrast, SspB-mScarlett exhibited strong polar localization and robust co-localization with SGP. If native SspB joins cytoplasmic SspB-mScarlett in being sequestered in stem cells, this could have the effect of reducing SspB-mediated T7RNAP-ssrA degradation in newborn factory cells. To test this hypothesis, we used Western Blotting to observe the level of T7RNAP-ssrA. We found that T7RNAP-ssrA levels were higher in MiST cultures than in the factory cell-only control and in cultures that employed xGP as the stem cell determinant (Supplementary Figure 3-D). These observations support the hypothesis that MiST^P^ factory cells gain a production boost from increased stability of T7RNAP-ssrA (Figure 4D).

**Figure 4.**
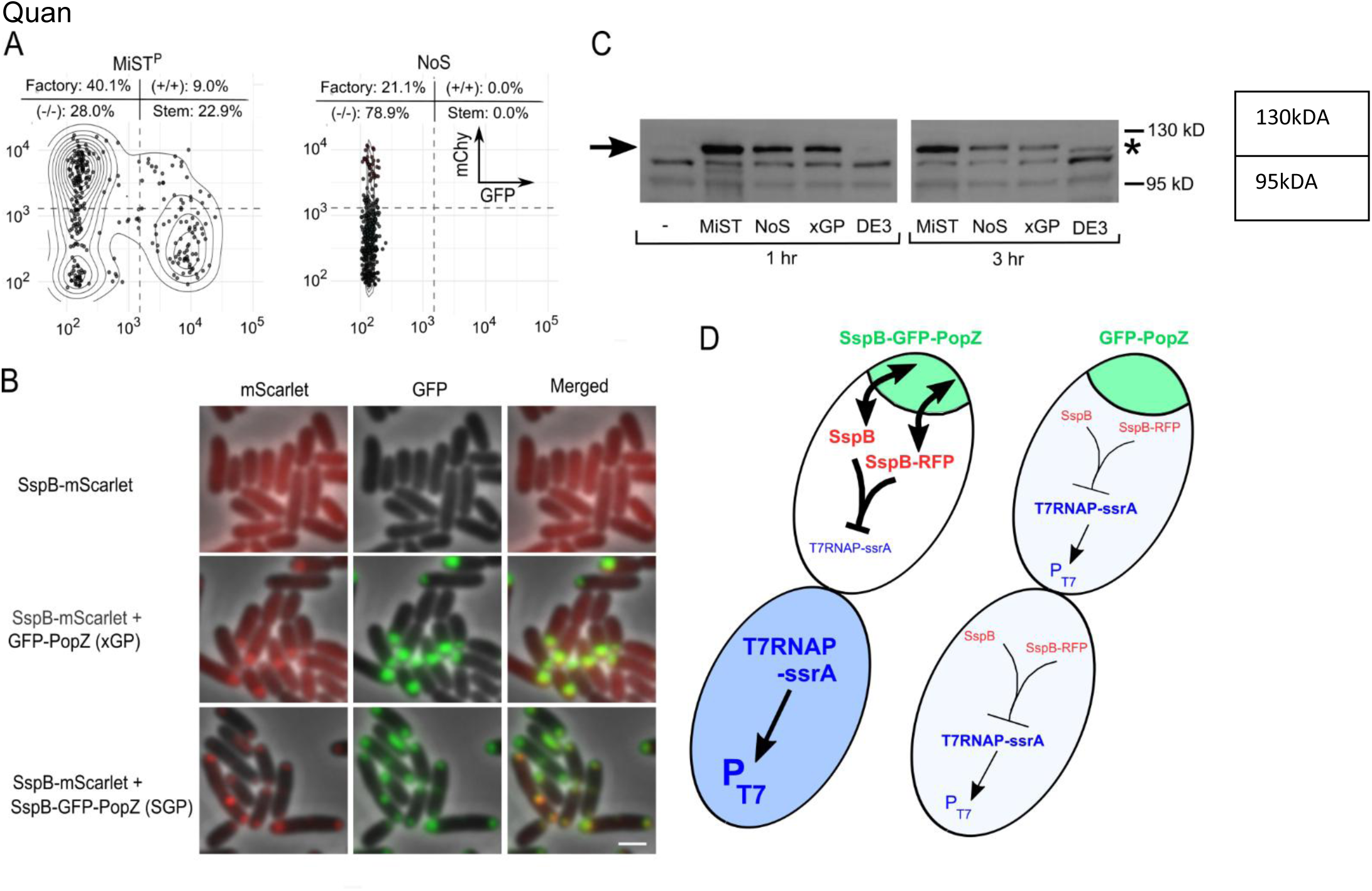
MiST stem cells support elevated target gene expression in factory cell progeny. (A) mCHY (Y-axis) and GFP (X-axis) fluorescence intensities in individual cells in stem cell-supported (MiST^P^) and factory cell-only (NoS) cultures 3 hours after the induction of factory cell differentiation. Arbitrary values (dotted lines) distinguish cell types. (B) Distribution of SspB-mScarlet (red) in control cells and in cells co-expressing SspB-GFP-PopZ (SGP) or GFP-PopZ (xGP) as stem cell determinants (green). Fluorescence images are overlayed on phase contrast images (gray). Scale bars 2µM. (C) T7 RNA Polymerase levels during product synthesis. Factory cell differentiation was induced in stem cell-supported MiST^P^ cultures (MiST) or in cultures expressing no stem cell determinant (NoS) or a truncated GFP-PopZ stem cell determinant (xGP) by incubation in 10 µM vanillate for 1 or 3 hours, and compared to MiST^P^ cultures not exposed to vanillate (-). T7RNAP expression was induced in conventional BL-21(DE3) cells with 200 µM IPTG. T7RNAP-ssrA (arrow) and T7RNAP (asterisk) were labeled by immunoblotting (arrow). (D) Model for T7RNAP-ssrA accumulation. In MiST cultures (left), polar sequestration of cytoplasmic SspB by the SGP stem cell determinant (green) results in an asymmetric distribution of T7RNAP-ssrA degradation machinery at cell division (red), resulting in stem cells that rapidly clear T7RNAP-ssrA and factory cells that rapidly accumulate T7RNAP-ssrA, stimulating target gene expression from *P_T7_*(blue). In a culture without SGP, cytoplasmic SspB is equally partitioned at cell division, leading to more T7RNAP-ssrA degradation in factory cells and consequently lower levels of expression from *P_T7_*.

### Chromosomally Integrated MiST Strains

A limitation of the aforementioned genetic circuitry is that it relies on the co-maintenance of three plasmids, which introduces genetic instability and imposes burdens associated with plasmid replication and segregation. To alleviate this issue, we used the SAGE (Serine recombinase-Assisted Genome Engineering ^55^ method to integrate some of the MiST genetic components into the *E. coli* chromosome. We inserted P_van_-T7RNAP-ssrA and the stem cell determinant P_BAD_-SGP at the HK022 phage integration site. This strain exhibited low SGP expression compared to our plasmid-based strains, likely because of the reduction in gene copy number, making it difficult to establish stem cells. We therefore reintroduced the plasmid-borne P_BAD_-SGP component of our system. To reduce the plasmid copy burden, we also moved the output cassette (either P_T7_-mChy alone or P_T7_-mChy with P_T7_-gp2) from the medium-copy pCDF plasmid backbone to low-copy pACYC, completing a revised version of the genetic circuitry, designated MiST^C1^ (Figure 5A).

**Figure 5.**
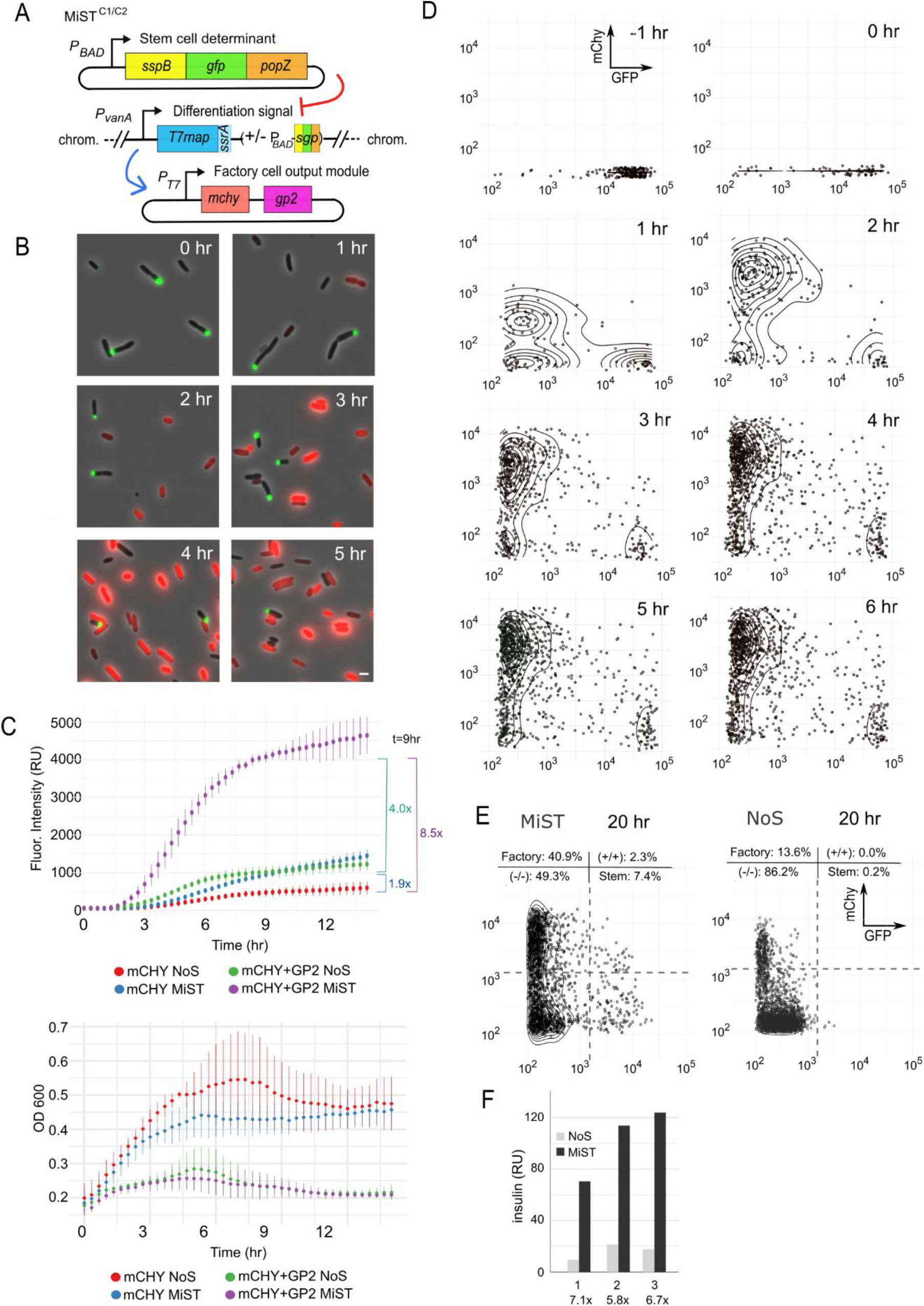
Genomic integration of MiST components provides more consistent performance and further boosts in target gene expression. (A) MiST^C1/C2^ genetic circuit, with the T7RNAP-ssrA cassette integrated into the chromosome, either together with (C1) or without (C2) a cassette for the SGP stem cell determinant. These strains also include the SGP stem cell determinant and the *P_T7_* output cassettes on separate plasmids. (B) Fluorescence images of MiST^C1^ cells after the induction of factory cell differentiation at t = 0 hr. SGP stem cell determinant (green polar foci) and factory cell product (red mCHY signal) are overlayed on a phase contrast image (gray). Scale bars 2µM. (C) mCHY fluorescence (upper panel) and OD600 (lower panel) in stem cell-supported MiST^C1^ cultures and factory cell-only control cultures, either with or without co-expression of gp2 in factory cells. Cells were grown in a microplate reader. Plots show average values from three independent experiments (bars: standard deviation). Average fold differences at 7 hr are denoted by brackets. (D) mCHY (Y-axis) and GFP (X-axis) fluorescence intensities in individual cells in stem cell-supported MiST^C1^ cultures, before and after inducing factory cell differentiation. (E) mChy and GFP fluorescence intensities in individual cells, as in D, except that stem cell-supported (MiST) and factory cell-only (NoS) cultures are compared after 20 hours of production and periodic culture dilution in fresh media. (F) Insulin produced by stem cell-supported MiST^C2^ cultures (MiST) and factory cell-only control cultures (NoS) after 24 hours of production, in three independent experiments, with fold differences indicated.

Sequential expression of the SCP stem cell determinant and T7RNAP-ssrA resulted in robust asymmetric cell division and clear differentiation of stem and factory cells (Figure 5B, Supplementary Movie M4). We evaluated mChy production and culture growth in microtiter plates, comparing MiST^C1^ cultures to factory cell-only controls (Figure 5C). At nine hours post-factory cell induction, MiST^C1^ cultures from the strain that did not include *gp2* produced 1.9-fold more mChy than factory cell-only controls. The total level of mChy production was highest in cells that co-expressed Gp2 with mChy. Here, stem cell-supported cultures accumulated 4.0x-fold more mChy than factory cell-only controls, and 8.5x-fold more mChy than cultures that did not co-express gp2 and did not have stem cell support. We followed the population dynamics of MiST^C1^ cultures in shake flask cultures by plotting the GFP and mChy fluorescence intensities of individual cells (Figure 5D). Compared to similar analyses of the three-plasmid MiST^P^ system (Supplementary Figure S3B), we observed a lower percentage of double-negative cells and an increase in the average mChy accumulation in factory cells (Supplementary Figure S4A). Together, these results indicate that the MiST^C1^ strain, with chromosomally integrated components, exhibits more robust cell differentiation and higher product yield.

### Demonstrating the functionality of MiST components

We built a series of control strains to test the functions of MiST control elements (Table 1). To test the influence of the SspB protease adaptor, we generated a chromosomal integrant that includes only P_VAN_-T7ssrA without an adjacent SGP gene. To this strain background, we added a P_T7_ output cassette and either plasmid-borne P_BAD_-SGP, driving expression of the normal stem cell determinant (a strain designated MiST^C2^), or plasmid-borne P_BAD_-xGP, driving expression of a stem cell determinant that is comprised of the first 13 amino acids of SspB fused to GFP-PopZ. While the MiST^C2^ strain exhibited robust asymmetric cell division and factory cell differentiation, the strain expressing the SspB-defective stem cell determinant was not clearly differentiated, with many GP-containing cells also positive for mChy expression (Supplementary Figure S4A, S4B). This indicates that the MiST stem cell determinant must include full-length SspB in order to achieve robust factory cell differentiation.

To test the influence of the ssrA protein degradation sequence, we generated a chromosomal integrant with a T7RNAP variant that lacks the SsrA tag. We attempted to co-transform this strain with the P_T7_-mChy-P_T7_-Gp2 output and P_BAD_-SGP plasmids, but never obtained viable colonies (Table 1). We hypothesized that low-level accumulation of T7RNAP and therefore leaky expression of Gp2 was responsible for the lack of viability, and tested this by asking if the strain could be co-transformed with pB-SGP and a P_T7_-mChy output cassette that lacks Gp2. This strain (designated MiST^X^) was viable, but exhibited leaky expression of mChy (Supplementary Figure S4C). We also found that the P_T7_-mChy-P_T7_-Gp2 output plasmid caused similar viability problems in a standard BL21(De3) strain, which harbors normal T7RNAP without an SsrA tag. Further, when we tested the P_T7_-mChy output cassette that lacks Gp2 in the BL21(De3) background, we found that many cells, including those expressing the SGP stem cell determinant, exhibited leaky expression of mChy (Supplementary Figure S4C).

From these experiments, we conclude that T7RNAP must be destabilized by tagging it with SsrA in order to prevent leaky Gp2 expression from MiST output plasmids.

### Assessing the Performance of chromosomally integrated MiST components

We monitored cell type differentiation and mChy production levels in MiST^C2^, the simplest and most widely applicable MiST strain background produced in this work (Figure 5E). After 20 hours of induction in shake flasks, MiST^C2^ cultures and factory-cell only controls reached stationary phase and included a substantial fraction of double-negative cells. The MiST^C2^ cultures, however, retained approximately 27 percent more factory cells, which accumulated nearly two-fold more mChy than factory cell-only controls, suggesting that stem cell support provides some protection against the proliferation of non-productive cheater mutants, even over extended times of production.

We also tested proinsulin production in MiST^C2^ strains (Figure 5F). Here, the pro-insulin coding sequence, tagged with a FLAG epitope, was placed into an output cassette that enabled co-expression of pro-insulin with mCherry and Gp2 (pCDF-Ins-mChy-Gp2). Stem cell-supported (MiST) and factory cell-only control cultures were grown in parallel in 50 mL shake flasks, and whole-cell lysates were collected after 20 hours of production and probed via Western blotting with an anti-FLAG antibody. MiST^C2^ cultures outperformed similar plasmid–based MiST^P^ strains (Figure 3E), accumulating an average of 6.5-fold more proinsulin than cultures that lacked stem cell support. These findings demonstrate the potential of MiST as a versatile and scalable platform for industrial bioproduction, particularly when the products are cytotoxic or otherwise inimical to cell growth.

## DISCUSSION

In this work, we merge the powerful gene expression capabilities of T7RNAP with the programmable cell differentiation capabilities of Microbial Stem Cell Technology (MiST).

The components of this genetic circuitry, which we name MiST2.0, are: i) an asymmetrically inherited cue, named SGP, which is passed into stem cells and excluded from factory cells; ii) an ssrA-tagged variant of T7RNAP that is degraded in SGP-containing stem cells and stable in factory cells; and iii) an expression output cassette that utilizes the T7RNAP-driven *P_T7_* promoter. We demonstrated that MiST2.0 cells can be induced to divide asymmetrically with respect to SGP, that SGP-containing stem cells remain mostly off for T7RNAP-driven gene expression, and that cells without SGP differentiate into factory cells in which T7RNAP drives gene expression from the *P_T7_* promoter.

In earlier reports, we and others have used stable macromolecular assemblies of PopZ as platforms for controlling cell behavior, as a cue for asymmetric cell division in *E. coli* and as a tool for controlling protein localization in eukaryotic cells^24^. The key determinant of these systems’ behaviors is PopZ functionalization. When we fused PopZ to an enzyme that cleaves cyclic-di-GMP, this generated a polarized signal that could be coupled to differential transcriptional output in stem cells and factory cells ^18^. While this system is capable of asymmetric cell division and cell differentiation, the transcriptional response is weak and incompatible with high-level expression systems typically used in biomanufacturing. Others have functionalized PopZ microdomains in *E. coli* as sites for the assembly of split-T7RNAP or a split-protease^20^. However, establishing cellular asymmetry in either of these systems relies upon coupling the activity of the reconstituted split-enzyme to the local abundance of a downstream protein that remains polarized on account of its slow diffusion rate, limiting the range of system outputs and their maximum expression levels. Another notable example was to use PopZ foci as gatekeepers for the accumulation of the metabolic intermediate shikimate in *E. coli*. Here, cells that inherited PopZ retained the ability to divert shikimate to anabolic pathways that supported cell growth, while their differentiated siblings lost this ability and accumulated shikimate as an end product. Although this system achieves modest increases in product titer over factory cell-only controls ^21^, its design is specific to the shikimate pathway.

MiST2.0 brings microbial stem cell technology substantially closer to the long-term goal of applying microbial stem cell technology to large-scale bio-manufacturing. Two significant advantages are that the factory cell output is driven by a highly active variant of T7RNAP, and the use of standard *P_T7_* output promoters makes the system compatible with widely used high-production chassis based on pET and similar T7RNAP-driven expression vectors. To demonstrate the usefulness of microbial stem cells in bio-manufacturing, we compared the production yields from MiST2.0 cultures to conventional cultures in which all cells are instructed to act as factory cells. We found that MiST cultures expressed up to 8.5-fold more target protein. Notably, we observed the highest increases in yield when product synthesis was coupled to cytotoxicity or the inhibition of cell growth. Since many bio-manufacturing operations are limited by the inhibitory effects of biosynthetic burden^2,56,57^, the context in which MiST2.0 provides the greatest benefit to production appears well-placed to address a major challenge for bio-industries.

One explanation for the increased yields in MiST2.0 cultures is that the system focuses cells’ gene expression machinery on T7RNAP target genes. MiST 2.0 incorporates the method developed by Stargardt *et al.,* wherein T7RNAP is co-expressed with a phage-derived RNA polymerase inhibitor peptide, called GP2, which specifically inhibits host cell RNAP but not T7RNAP^46,53^. This method is reported to provide 2.7 to 3.4-fold higher target protein yield, and we observed a similar outcome (1.9-fold increase) in analogous experiments, where GP2 was co-expressed with T7RNAP in non-MiST factory cell-only cultures.

An important difference between MiST2.0 and conventional factory cell-only production methods is that the former includes a stem cell sub-population that is protected from the growth-inhibiting effects of GP2 and other engineered products. Our time-lapse microscopy experiments showed that MiST stem cells continued to elongate and divide while their factory cell progeny experienced growth arrest. We also observed that MiST cultures consistently produced higher optical densities than factory cell-only controls, although the difference was usually less than 2-fold and not as large as we expected. It is possible that stem cells experience leaky GP2 expression, limiting their growth. It is also possible that stem cell-derived population growth is partially overshadowed by the rapid growth of “cheater mutants”, such as factories that have lost the ability to produce GP2. Consistent with this, we observed a progressive rise in the fraction of “double-negative” cells, which expressed neither mCherry nor GFP, in long-term growth experiments. This problem might be alleviated by stabilizing the GP2 output cassette as a chromosomal integration.

The modest increase in optical densities of MiST cultures compared to factory cell-only controls is not sufficient to explain the multi-fold increases observed for target protein synthesis. Instead, most of the yield increase can be explained by the fact that MiST-generated factory cells express more target protein than cells generated under factory cell-only conditions. We found that the stem cell determinant in MiST cultures sequesters native SspB, and this is probably the reason for the increased stability of T7RNAP-ssrA in factory cells in MiST cultures and higher levels of T7RNAP-induced target gene expression. Overall, we find that three different processes work in parallel to support higher yields in MiST cultures: i) an underlying stem cell population supports continued generation of new factory cells; ii) GP2 focuses factory cell resources on T7RNAP-driven gene expression; and iii) SspB sequestration in stem cells allows higher T7RNAP levels in factory cells.

Another feature of the MiST2.0 system that elevates its potential for bio-manufacturing is that the polar sequestration mechanisms that support T7RNAP activity in factory cells also act to reduce leaky T7RNAP expression in stem cells. We found that this provides protection to MiST2.0 strains carrying plasmids with toxic output cassettes. Whereas the common T7RNAP expression strain BL21(De3) cannot be transformed with the *P_T7_-gp2* output plasmid, probably because leaky expression of GP2 inhibits cell growth, pre-induction cultures of MiST2.0 strains carrying the same plasmid exhibited robust growth and fluorescence microscopy confirmed that nearly all cells contained the plasmid. We also found that *P_T7_-gp2* output cassette cannot be carried by a MiST strain if T7RNAP lacks an ssrA tag, supporting the idea that stem cells are protected by the degradation of T7RNAP. We conclude that MiST2.0 is particularly well-suited for the production of highly toxic products that impose counterselection pressures even in the pre-induced state.

The concept of incorporating stem-cell-like activities into microbes has been explored by multiple groups, with the goal of increasing genetic stability of *E. coli* ^58^ and yeast ^59^ during product synthesis. Although the technical approaches are substantially different than MiST2.0, they highlight innovative growth in this field and the competitive landscape among technologies that seek to leverage the advantages of multicellularity. In this work, we demonstrate a mechanism for programming bacterial cell differentiation on the basis of T7RNAP activation. Given the widespread use of T7RNAP for diving product synthesis in biomanufacturing^60^, we propose MiST2.0 as a readily adaptable platform for leveraging the advantages of stem cells for high-titer yields during microbial biofermentations.

## Acknowledgments

The authors thank Dr. Adam Guss of Oak Ridge National Lab for providing advice and genetic components. This work was supported by Small Business Innovation Phase I grants to AsimicA Inc. and GB from the National Institute of Health (1R41GM137710) and the National Science Foundation (2222602), to AsimicA from the Wyoming Business Council, and to N.M. under the NSF Activate Fellowship Program.

---

**Supplementary Table 1.**
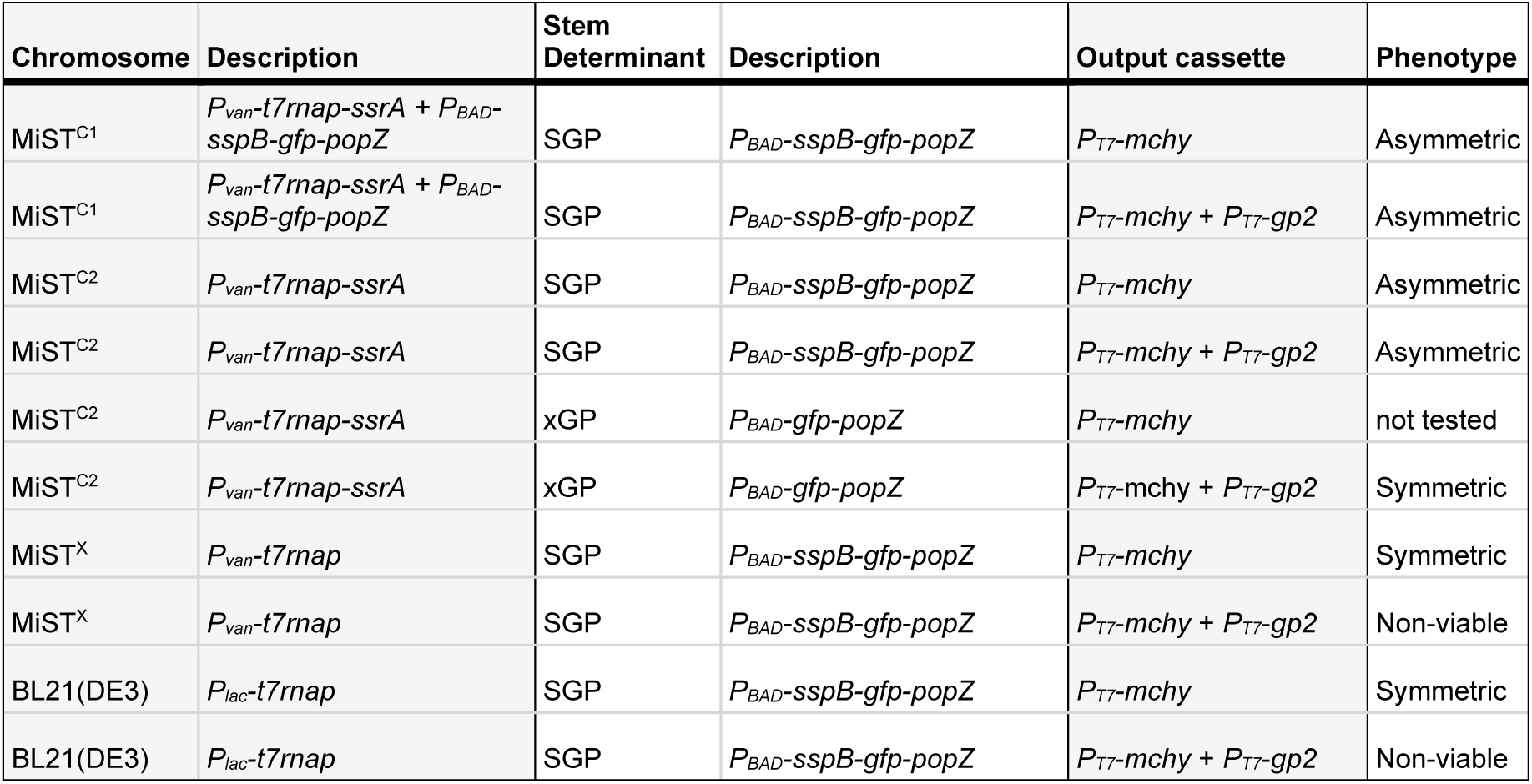
Functionality of MiST components in different strain backgrounds.

Table 1. Summary of phenotype characteristics of MiST and control strains carrying various output cassettes.

## Supplementary Figures

**Supplementary Figure 1.**
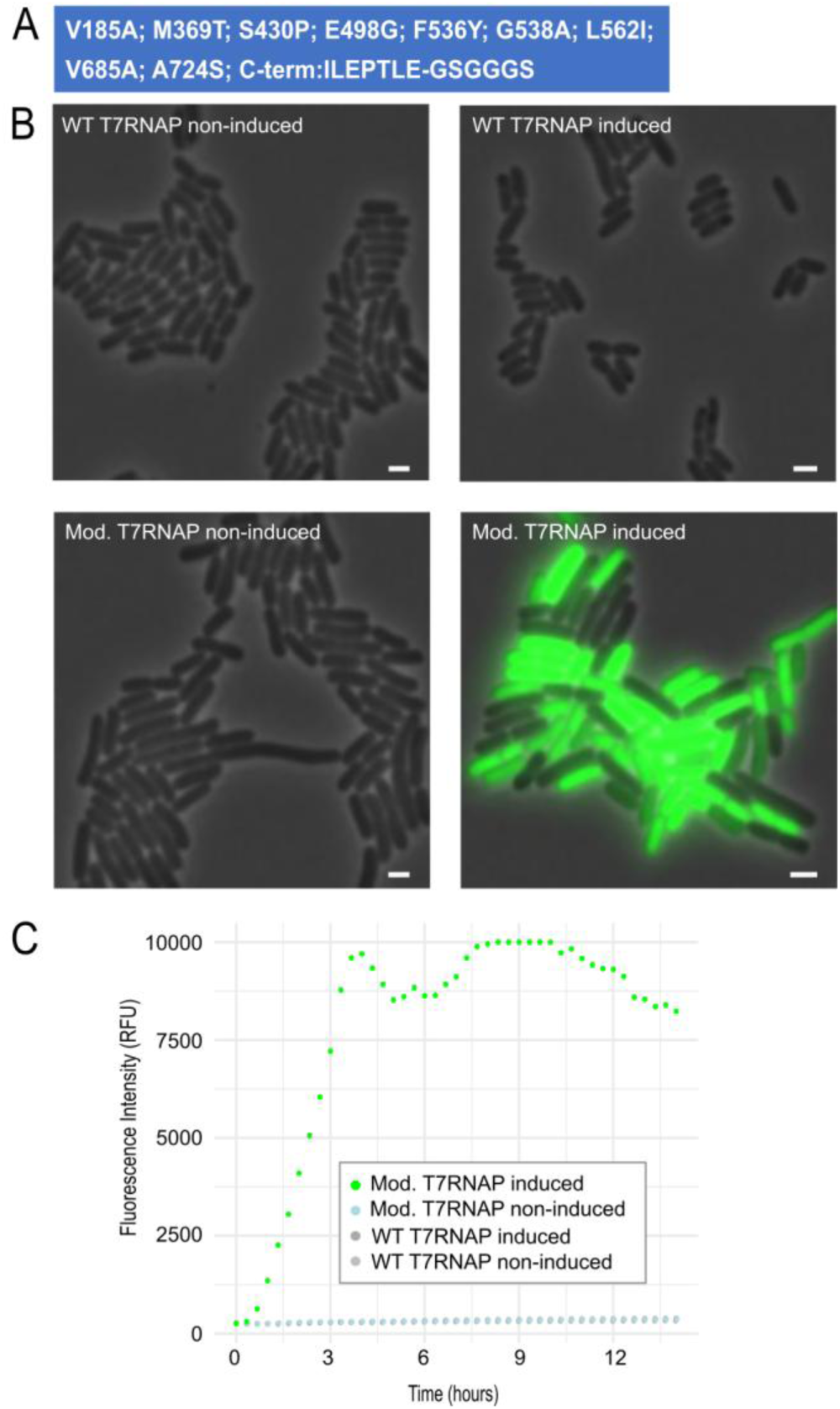
Functionalization of T7RNAP-ssrA. (A) Amino acid coding sequence changes introduced into Modified T7RNAP. (B) Induction of target gene expression by Wildtype (WT) or Modified (Mod) T7RNAP-ssrA. *E. coli* strains bearing the T7RNAP-ssrA variants under control of the *P_vanA_* promoter on plasmid pBAD were co-transformed with plasmid pACYC-Duet carrying the *gfp* target gene was under the *P_T7_* promoter. Cultures were uninduced (left) or induced with 50 µM vanillate and 50 µM IPTG for 2 hours prior to observation by fluorescence microscopy. GFP fluorescence signal (green) is overlayed on phase contrast (gray). Scale bars 2µM. (C) Strains from B were induced as above and grown in microplates for the indicated time (X-axis) with periodic measurement of GFP fluorescence (Y-axis).

**Supplementary Figure 2.**
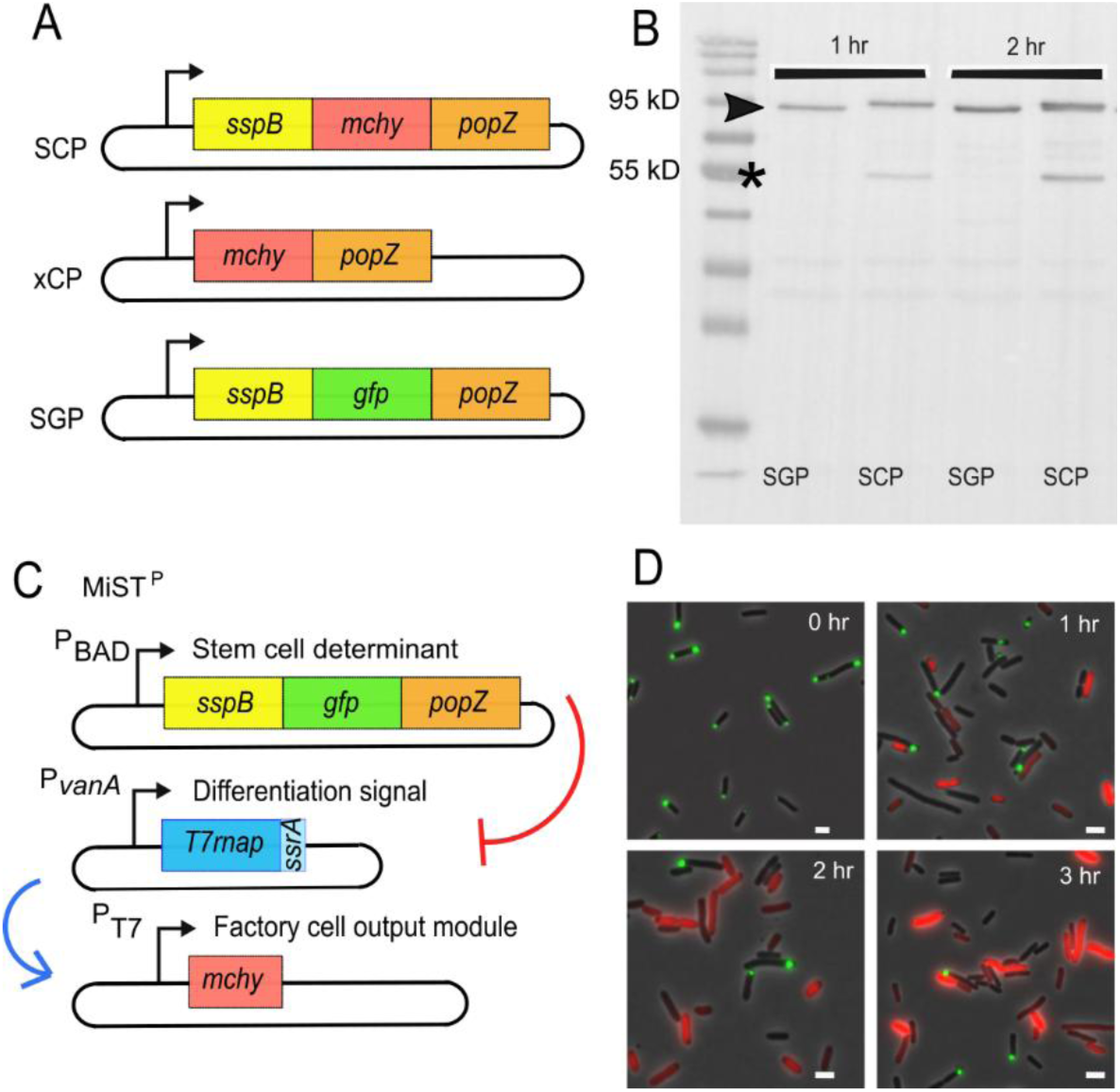
Modifications to stem cell determinant. (A) Gene diagram of SspB-mChy-PopZ (SCP), mChy-PopZ (xCP), and SspB-GFP-PopZ (SGP) stem cell determinants. (B) Western blot, using anti-PopZ primary antibody, of cultures expressing stem cell determinants. Full-length and truncated products are indicated by arrow and asterisk, respectively. (C) Genetic circuit of MiST^P^ with SGP stem cell determinant and mCHY expression as output. (D) Fluorescence images of MiST^P^ cells described in C after the induction of factory cell differentiation at t = 0 hr. SGP stem cell determinant (green polar foci) and factory cell product (red mCHY signal) are overlayed on a phase contrast image (gray). SGP expression was induced with 0.005% arabinose and T7RNAPssrA activity was induced with 15 µM vanillate and 15 µM IPTG. Scale bars 2µM.

**Supplementary Figure 3.**
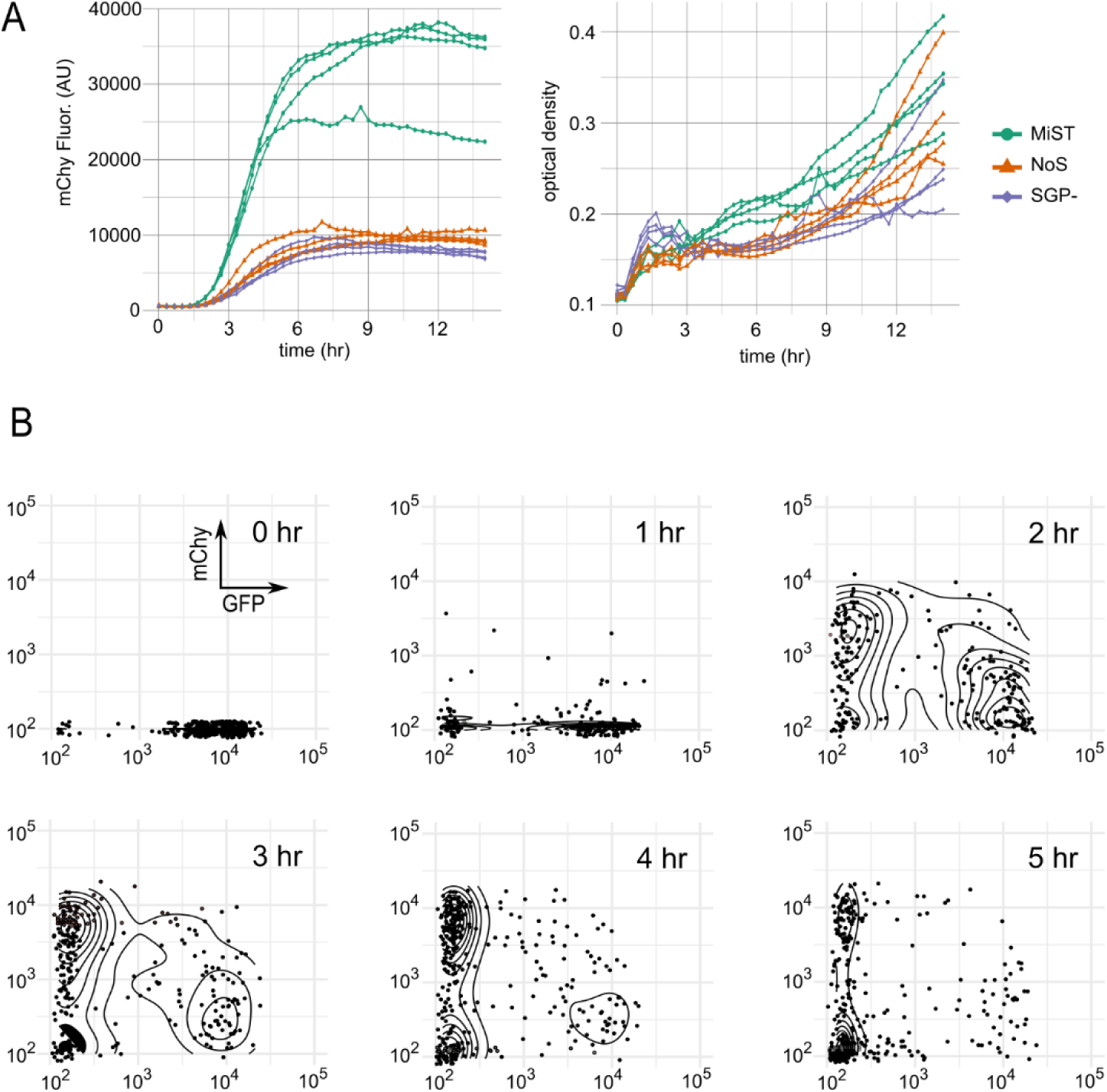
Stem cell-supported (MiST) cultures divide asymmetrically and express higher levels of target genes than factory cell-only cultures. (A) Observations of MiST^P^ cultures (MiST, green) and control cultures, either the same strain cultured under factory cell-only conditions (NoS, orange), or a strain with no SGP stem cell determinant and induced under MiST conditions (SGP-, purple). To assess clonal variation, each trial represents data from an independently isolated colony following co-transformation of plasmids containing system components. Cultures were grown in a microplate reader for the indicated time (X-axis), and mCHY fluorescence (Y-axis, left panel) and optical density (Y-axis, right panel) were collected over periodic intervals. (B) mCHY (Y-axis) and GFP (X-axis) fluorescence intensities of individual cells in stem cell-supported MiST^P^ cultures expressing mCHY and GP2 as factory cell outputs, with 0 hr as the time of factory cell induction.

**Supplementary Figure 4.**
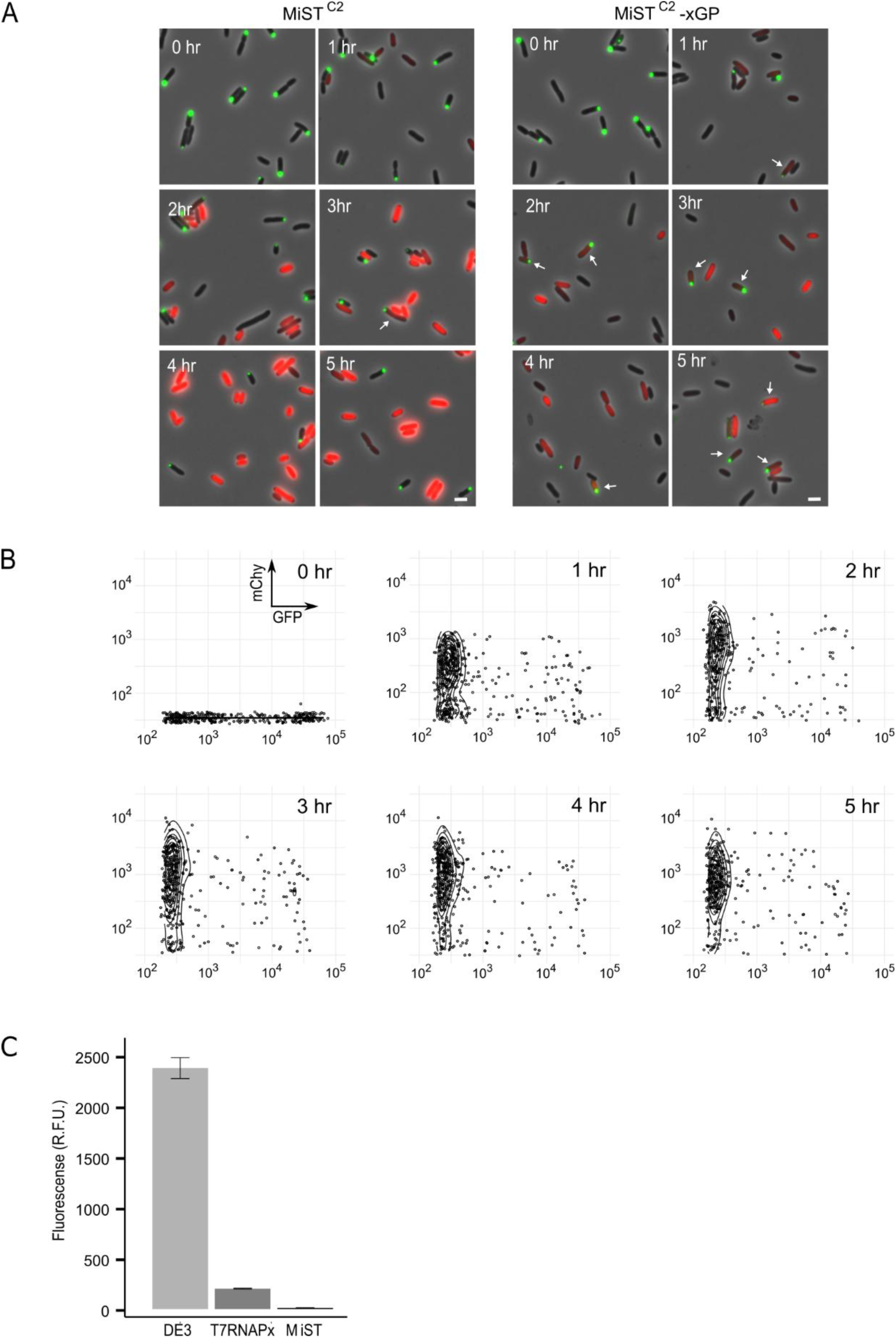
Consequences of expressing a non-functional stem cell determinant. (A) Fluorescence images of MiST cultures expressing a normal SGP stem cell determinant (MiST^C2^, left panels) or a truncated xGP stem cell determinant (MiST^C2^-xGP), with 0 hr as the time of factory cell induction. The stem cell determinant (green polar foci) and factory cell product (red mCHY signal with gp2, not visible) are overlayed on a phase contrast image (gray). Arrows indicate double-positive cells that express both mCHY and GFP. Scale bars 2µM. (B) mCHY (Y-axis) and GFP (X-axis) fluorescence intensities of individual cells in a MiST^C2^-xGP culture, with 0 hr as the time of factory cell induction. (C) Comparison of leaky target gene expression. A conventional BL21(DE3) strain (DE3, left), a MiST^C2^ strain expressing chromosomally integrated T7RNAP-ssrA (MiST, right), and a modified MiST strain expressing chromosomally integrated T7RNAP without a ssrA tag (T7RNAPx, center), all carrying plasmid-borne *P_T7_*-mCHY (without GP2) as an output cassette, were grown to stationary phase without any chemical inducers and their mCHY fluorescence intensity (Y-axis) was measured using a microplate reader.

## Supplementary Materials

**Supplementary Table 2.**
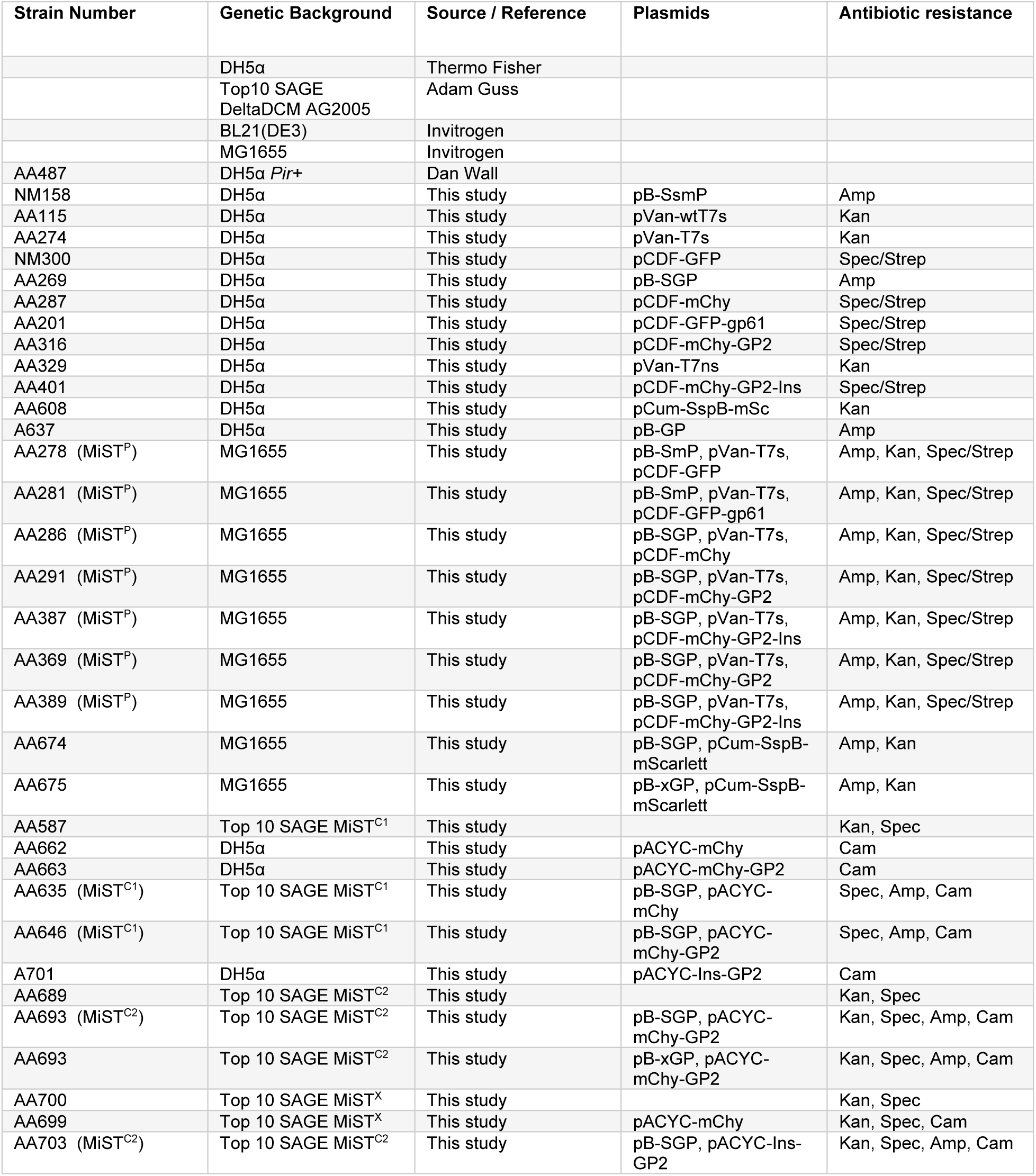
list of strains used in this study.

**Supplementary Table 3.**
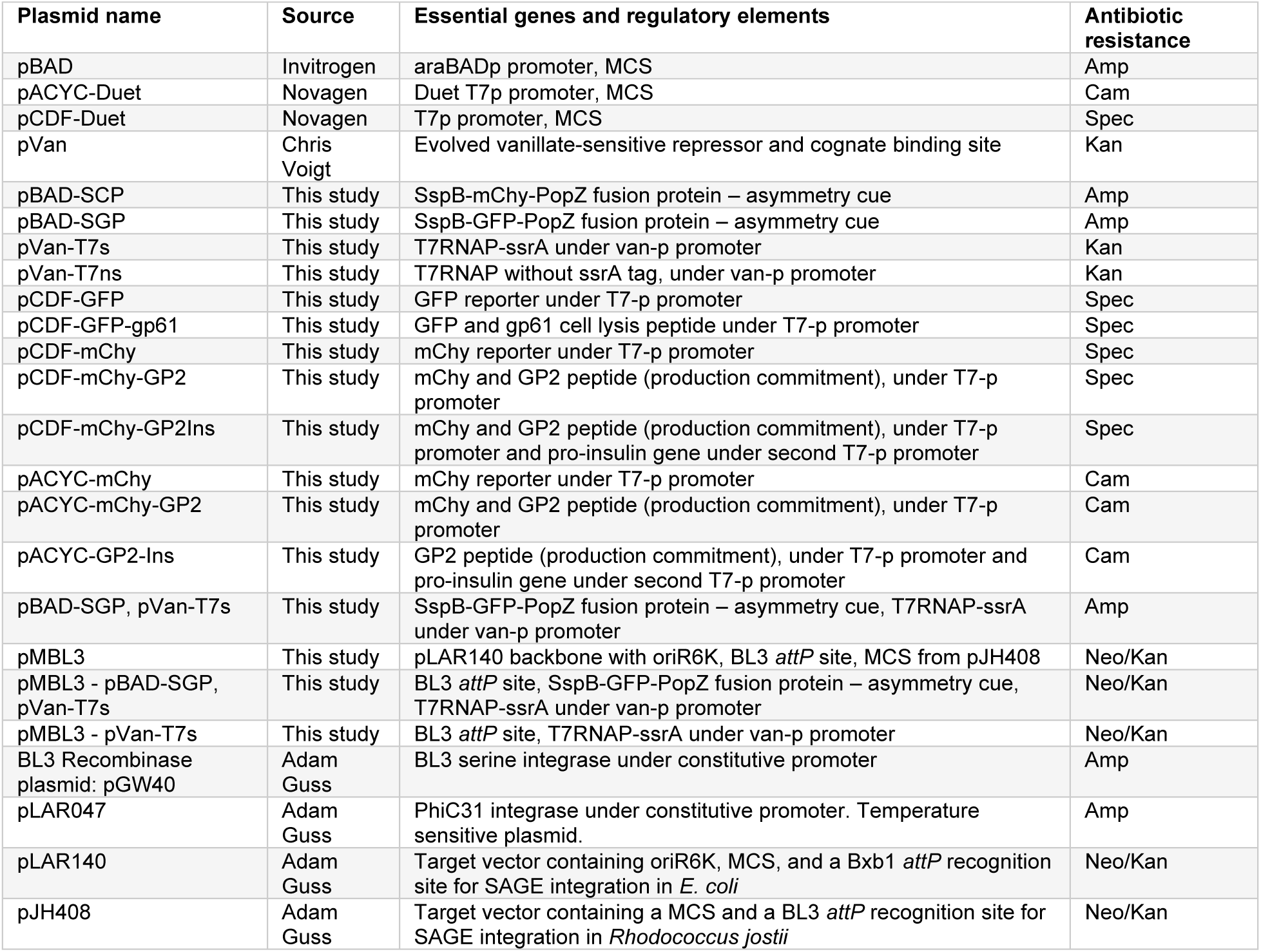
list of DNA plasmids used in this study.

## Materials and Methods

### Plasmid Construction

Plasmids were constructed by Isothermal Cloning (ITC) using the NEBuilder^®^ HiFi DNA Assembly Master Mix (New England Biolabs **Lot #10238675**), and where indicated, chemically synthesized DNA fragments (Integrated DNA Technologies gBlocks Gene Fragments). Plasmid names and key features are summarized in Supplementary Table 2.

### Polar degradation hub/asymmetry cues

To create pB-SCP, the sspB-mChy-PopZ fusion protein was cloned into a pBAD vector under the arabinose-inducible araBAD promoter using PCR amplification of individual genes with 9-amino acid linkers as primer extensions, followed by isothermal cloning (NEBuilder^®^ HiFi DNA Assembly Master Mix). The resulting plasmid was referred to as pB-SCP. To create pB-SGP, the msfGFP sequence was amplified via PCR and inserted via ITC into a PCR amplification product of pB-SCP that excluded the mCHY sequence.

### T7RNAP plasmids

Initially, the T7RNAP gene was amplified from the chromosome of the BL21(DE3) strain with the ssrA tag from *E. coli* (AANDENYALAA) added to the C-terminal end using PCR primer extension, and was inserted under the vanillate-inducible van-p promoter in the pVan plasmid ^61^, creating plasmid pVan-wtT7s. A mutated version of T7RNAP-ssrA (T7s-mut) was chemically synthesized to incorporate the set of mutations (see Supplementary Figure 1A) and cloned under the van-p promoter in the same manner as the wild-type counterpart, resulting in the pVan-T7s plasmid.

### Target gene plasmids

A fluorescent reporter marking differentiating gene expression in Factory cells was initially cloned in the pCDF-Duet vector under the T7-p promoter to facilitate T7RNAP-mediated expression. The monomeric superfolder GFP (msfGFP) was used as a reporter, resulting in the pCDF-GFP plasmid. Eventually, this was replaced with the mChy reporter to complement the redesigned polar asymmetry cue marked with GFP (pCDF-mChy). The DNA coding sequence of gp61 cell-wall lysis peptide was chemically synthesized and cloned downstream of the GFP reporter as a polycistronic gene in the pCDF-GFP plasmid, resulting in pCDF-GFP-gp61.

The DNA coding sequence of GP2 growth-arresting peptide was chemically synthesized and cloned downstream of the mChy reporter as a polycistronic gene in the pCDF-mChy plasmid, resulting in pCDF-mChy-GP2.

The DNA coding sequence of pre-pro-insulin gene was chemically synthesized and cloned under the second T7-p promoter in pCDF-Duet vector.

For chromosomally integrated MiST strains, the target genes were transferred from reporter plasmid pCDF-Duet to pACYC-Duet via PCR amplification and ITC cloning.

### Chromosomal Integration and Vector Backbone Excision

To create a target plasmid compatible with BL3 serine integrase, we PCR-amplified a region including a multiple cloning site (MCS) and a BL3 *attP* recognition site from plasmid pJH408 and a region including the entirety of pLAR140 with the exception of its MCS and the Bxb1 *attP* recognition site from plasmid pLAR140, then stitched the fragments together by isothermal cloning to make plasmid pMBL3. For SAGE (Serine Recombinase-Assisted Genome Engineering)-mediated genomic integrations, the Top10-SAGE target strain background contained a chromosomal landing pad site with an array of *attB* recognition sites for 14 serine integrases ^55^. DNA sequences intended for integration were first cloned into a neo/kan resistant target plasmid between the *attP* recognition sites for one of the 14 serine integrases. This plasmid has an oriR6K, which could only replicate in a Pir+ strain, and was co-transformed with a temperature-sensitive accessory plasmid carrying the cognate serine integrase into the Top10-SAGE target strain. Subsequent recovery at 37°C with kanamycin selected *E. coli* clones with integrated target genetic material and cured carrier plasmid. The same method was used to generate MiST^C1^ and MiST^C2^ strains.

To excise the integrated oriR6K-Neo/Kan backbone from the chromosome, PhiC31 recognition sites flanking the Neo/Kan markers within integrated DNA sequences, allowed for removal of the selective marker ^62^. The temperature sensitive plasmid containing PhiC31 (pLAR047) was transformed into the strain at 30°C then incubated overnight at 37°C to cure pLAR047. The culture was then diluted and plated out for single colonies on antibiotic-free plates. 25 colonies were then screened for the loss of both pLAR047 and the oriR6k-neo/kan backbone by spot plating the colonies on each antibiotic.

### Cell Culture

Strains were grown in Luria-Bertani medium (Miller formulation) and, where appropriate, with the following antibiotics: ampicillin (50 µg/ml), kanamycin (30 µg/ml), spectinomycin (50 µg/ml), chloramphenicol (20 µg/ml).

To prepare MiST and control strains for product synthesis trials, overnight cultures were diluted 100-fold in two 4 mL culture tubes and incubated for two hours in a tissue culture rotator (for strains carrying the GP2 peptide on the production plasmid, this incubation was extended to three hours). MiST cultures were then induced with 0.005% arabinose for one hour to express SspB-mChy/GFP-PopZ stem-cell determinant and asymmetry cue, and arabinose was omitted for factory cell-only control cultures. Next, 4 mL of culture was transferred to Eppendorf tubes, centrifuged at 9000 rpm for 2 minutes, then resuspended in LB medium. This was followed by two additional washes. Washed cells were then inoculated into 20 mL LB in 125ml Erlenmeyer flasks and incubated for one hour at 37°C with shaking at 250 rpm. At this stage, MiST cultures were determined to have returned to exponential growth and were primarily composed of stem cells with unipolar PopZ foci, marking the starting point for outcome expression experiments (zero time point, T0). The expression of output genes in the production plasmid (fluorescent markers, bacteriophage peptides, or pro-insulin) was induced by adding 2.5–15 μM vanillate and 2.5–15 μM IPTG (optimized for each strain). Aliquots of growing cultures were collected hourly for subsequent analyses. For microplate reader assays, cultures at the T0 point, prepared as described above, were transferred to sterile polystyrene 96-well plates with lids (100 μL per well). Plates were incubated in a Biotek Synergy H2 plate reader for 14 hours at 37°C with shaking, and fluorescence and optical density readings were obtained at 20 minute intervals.

### Fluorescence Microscopy and Image Analysis

Cells were immobilized on 1% agarose pads. Live-cell imaging was performed using a Zeiss Axio Imager Z2 epifluorescence microscope equipped with a Hamamatsu Orca-Flash4.0 sCMOS camera and a Plan-Apochromat 100x/1.46 Oil Ph3 objective. Zeiss filter sets 38HE and 63HE were used to acquire fluorescent images of msfGFP and mCherry, respectively. For time-lapse microscopy experiments, cells were immobilized on 1.2% low-melting-point agarose pads (Ultra PURE LMP agarose, GIBCO BRL, Life Technologies Inc.). Fluorescent images were taken with the same settings as described above, with image capture at 20-minute intervals.

Images were collected and processed with Zen Blue software. For image quantitation, fluorescence micrographs were split into three channels (phase contrast, GFP, mChy) and exported as TIFF files. Stack images were re-created using Fiji (ImageJ) software. Detection of bacterial cells was performed using the MicrobeJ plugin. The maximum fluorescence signal for the stem cell marker and the mean signal for expressed fluorescent output were collected for individual cells. Data plots were constructed using R software.

### Western Blotting (WB)

Cell lysates were prepared by mixing 100 μL cell cultures with sample buffer (NEB) and boiling at >90°C for 10 minutes, then resolved by SDS-PAGE and transferred to PVDF membranes according to the manufacturer’s instructions. Membranes were incubated with primary antibodies (anti-FLAG (Proteintech, Cat No. 20543-1-AP, polyclonal, 1:5000 concentration), anti-PopZ (rabbit polyclonal, 1:2,000), or anti-T7RNAP (Proteintech, Cat No. 29943-1-AP, polyclonal, 1:5000 concentration)) for one hour, washed three times in PBS-Tween, incubated with secondary antibodies (anti-rabbit conjugated with HRP, Jackson Immunoresearch), and washed again three times. Membranes were then incubated with a chemiluminescence substrate kit (ThermoFisher) for five minutes and used for signal detection and image collection (Bio-Rad gel documentation system).

## REFERENCES

1. Glick, B. R. Metabolic Load and Heterologous Gene Expression. Biotechnol Adv 13, 247–261 (1995).

2. Rajacharya, G. H., Sharma, A. & Yazdani, S. S. Proteomics and metabolic burden analysis to understand the impact of recombinant protein production in E. coli. Sci Rep 14, (2024).

3. Straathof, A. J. J. Modelling of end-product inhibition in fermentation. Biochemical Engineering Journal vol. 191 Preprint at 10.1016/j.bej.2022.108796 (2023).

4. Rugbjerg, P., Myling-Petersen, N., Porse, A., Sarup-Lytzen, K. & Sommer, M. O. A. Diverse genetic error modes constrain large-scale bio-based production. Nat Commun 9, (2018).

5. Hoffmann, F. & Rinas, U. Stress induced by recombinant protein production in Escherichia coli. Advances in biochemical engineering/biotechnology vol. 89 73–92 Preprint at 10.1007/b93994 (2004).

6. Alper, H. S. & Avalos, J. L. Metabolic pathway engineering. Synthetic and Systems Biotechnology vol. 3 1–2 Preprint at 10.1016/j.synbio.2018.01.002 (2018).

7. Ding, S. et al. From reactants to products: computational methods for biosynthetic pathway design. Synthetic and Systems Biotechnology vol. 10 1038–1049 Preprint at 10.1016/j.synbio.2025.05.005 (2025).

8. Orsi, E., Schada von Borzyskowski, L., Noack, S., Nikel, P. I. & Lindner, S. N. Automated in vivo enzyme engineering accelerates biocatalyst optimization. Nat Commun 15, (2024).

9. Cravens, A., Payne, J. & Smolke, C. D. Synthetic biology strategies for microbial biosynthesis of plant natural products. Nature Communications vol. 10 Preprint at 10.1038/s41467-019-09848-w (2019).

10. Chae, T. U., Choi, S. Y., Kim, J. W., Ko, Y. S. & Lee, S. Y. Recent advances in systems metabolic engineering tools and strategies. Current Opinion in Biotechnology vol. 47 67–82 Preprint at 10.1016/j.copbio.2017.06.007 (2017).

11. Das, M., Patra, P. & Ghosh, A. Metabolic engineering for enhancing microbial biosynthesis of advanced biofuels. Renewable and Sustainable Energy Reviews vol. 119 Preprint at 10.1016/j.rser.2019.109562 (2020).

12. Xu, P. Production of chemicals using dynamic control of metabolic fluxes. Current Opinion in Biotechnology vol. 53 12–19 Preprint at 10.1016/j.copbio.2017.10.009 (2018).

13. Raman, S., Rogers, J. K., Taylor, N. D. & Church, G. M. Evolution-guided optimization of biosynthetic pathways. Proc Natl Acad Sci U S A 111, 17803–17808 (2014).

14. Lechner, A., Brunk, E. & Keasling, J. D. The need for integrated approaches in metabolic engineering. Cold Spring Harbor Perspectives in Biology vol. 8 Preprint at 10.1101/cshperspect.a023903 (2016).

15. Du, Y. H. et al. Optimization and Scale-Up of Fermentation Processes Driven by Models. Bioengineering vol. 9 Preprint at 10.3390/bioengineering9090473 (2022).

16. Boodhoo, K. V. K., Flickinger, M. C., Woodley, J. M. & Emanuelsson, E. A. C. Bioprocess intensification: A route to efficient and sustainable biocatalytic transformations for the future. Chemical Engineering and Processing - Process Intensification vol. 172 Preprint at 10.1016/j.cep.2022.108793 (2022).

17. Wang, Z. Z. et al. Fermentation design and process optimization strategy based on machine learning. BioDesign Research vol. 7 Preprint at 10.1016/j.bidere.2025.100002 (2025).

18. Mushnikov, N. V., Fomicheva, A., Gomelsky, M. & Bowman, G. R. Inducible asymmetric cell division and cell differentiation in a bacterium. Nat Chem Biol 15, 925–931 (2019).

19. Bowman, G., Mushnikov, N. & Gomelsky, M. 11525117 Issued Patent: Microbial Stem Cell Technology.

20. Lin, D. W. et al. Construction of intracellular asymmetry and asymmetric division in Escherichia coli. Nat Commun 12, (2021).

21. Ding, Q. et al. Engineering Escherichia coli asymmetry distribution-based synthetic consortium for shikimate production. Biotechnol Bioeng 119, 3230–3240 (2022).

22. Bowman, G. R. et al. Caulobacter PopZ forms a polar subdomain dictating sequential changes in pole composition and function. Mol Microbiol 76, 173–189 (2010).

23. Ehrle, H. M. et al. Polar organizing protein PopZ is required for chromosome segregation in Agrobacterium tumefaciens. J Bacteriol 199, (2017).

24. Lasker, K. et al. The material properties of a bacterial-derived biomolecular condensate tune biological function in natural and synthetic systems. Nat Commun 13, (2022).

25. Bowman, G. R. et al. Oligomerization and higher-order assembly contribute to sub-cellular localization of a bacterial scaffold. Mol Microbiol 90, 776–795 (2013).

26. Ebersbach, G., Briegel, A., Jensen, G. J. & Jacobs-Wagner, C. A Self-Associating Protein Critical for Chromosome Attachment, Division, and Polar Organization in Caulobacter. Cell 134, 956–968 (2008).

27. Studier, F. W. & Moffattf, B. A. Use of Bacteriophage T7 RNA Polymerase to Direct Selective High-level Expression of Cloned Genes. J. MoZ. Biol 189, 113–130 (1986).

28. Kushwaha, M. & Salis, H. M. A portable expression resource for engineering cross-species genetic circuits and pathways. Nat Commun 6, (2015).

29. Brunsehwig, E. & Darzins, A. A Two-Component T7 System for the Overexpression of Genes in Pseudomonas Aeruginosa. (1992).

30. Effendi, S. S. W. & Ng, I. S. Innovations, Challenges and Future Directions of T7RNA Polymerase in Microbial Cell Factories. ACS Synthetic Biology Preprint at 10.1021/acssynbio.5c00139 (2025).

31. Du, F. et al. Regulating the T7 RNA polymerase expression in E. coli BL21 (DE3) to provide more host options for recombinant protein production. Microb Cell Fact 20, (2021).

32. Tabor, S. Expression Using the T7 RNA Polymerase/Promoter System . Curr Protoc Mol Biol 11, (1990).

33. Heyde, S. A. H. & Nørholm, M. H. H. Tailoring the evolution of BL21(DE3) uncovers a key role for RNA stability in gene expression toxicity. Commun Biol 4, (2021).

34. Robichon, C., Luo, J., Causey, T. B., Benner, J. S. & Samuelson, J. C. Engineering Escherichia coli BL21(DE3) derivative strains to minimize E. coli Protein contamination after purification by immobilized metal affinity chromatography. Appl Environ Microbiol 77, 4634–4646 (2011).

35. Dubendorfft, J. W. & Studier, F. W. Controlling Basal Expression in an Inducible T7 Expression System by Blocking the Target T7 Promoter with lac Repressor. J. Mol. Biol 219, 45–59 (1991).

36. Sanchez, A., Osborne, M. L., Friedman, L. J., Kondev, J. & Gelles, J. Mechanism of transcriptional repression at a bacterial promoter by analysis of single molecules. EMBO Journal 30, 3940–3946 (2011).

37. Shilling, P. J., et al. Improved designs for pET expression plasmids increase protein production yield in Escherichia coli. Commun Biol 3, (2020).

38. Rosenberg, A. H. et al. Vectors for selective expression of cloned DNAs by T7 RNA polymerase. Gene 56, 125–135 (1987).

39. Du, F. et al. Regulating the T7 RNA polymerase expression in E. coli BL21 (DE3) to provide more host options for recombinant protein production. Microb Cell Fact 20, (2021).

40. Iost, I., Guillerez, J. & Dreyfus, M. NOTES Bacteriophage T7 RNA Polymerase Travels Far Ahead of Ribosomes In Vivo. J Bacteriol 174, 619–622 (1992).

41. Shis, D. L. & Bennett, M. R. Synthetic biology: the many facets of T7 RNA polymerase. Mol Syst Biol 10, (2014).

42. Wagner, S. et al. Consequences of membrane protein overexpression in Escherichia coli. Molecular and Cellular Proteomics 6, 1527–1550 (2007).

43. Kim, S. K., Lee, D. H., Kim, O. C., Kim, J. F. & Yoon, S. H. Tunable Control of an Escherichia coli Expression System for the Overproduction of Membrane Proteins by Titrated Expression of a Mutant lac Repressor. ACS Synth Biol 6, 1766–1773 (2017).

44. Conrad, T., Plumbom, I., Alcobendas, M., Vidal, R. & Sauer, S. Maximizing transcription of nucleic acids with efficient T7 promoters. Commun Biol 3, (2020).

45. Li, S. et al. Enhanced protein and biochemical production using CRISPRi-based growth switches. Metab Eng 38, 274–284 (2016).

46. Stargardt, P., Feuchtenhofer, L., Cserjan-Puschmann, M., Striedner, G. & Mairhofer, J. Bacteriophage Inspired Growth-Decoupled Recombinant Protein Production in Escherichia coli. ACS Synth Biol 9, 1336–1348 (2020).

47. Pu, J., Disare, M. & Dickinson, B. C. Evolution of C-Terminal Modification Tolerance in Full-Length and Split T7 RNA Polymerase Biosensors. ChemBioChem 20, 1547–1553 (2019).

48. Gottesman, S., Roche, E., Zhou, Y. & Sauer, R. T. The ClpXP and ClpAP proteases degrade proteins with carboxy-terminal peptide tails added by the SsrA-tagging system. Genes Dev 1338–1347 (1998).

49. Lies, M. & Maurizi, M. R. Turnover of endogenous SsrA-tagged proteins mediated by ATP-dependent proteases in Escherichia coli. Journal of Biological Chemistry 283, 22918–22929 (2008).

50. Dougan, D. A., Weber-Ban, E. & Bukau, B. Targeted Delivery of an ssrA-Tagged Substrate by the Adaptor Protein SspB to Its Cognate AAA Protein ClpX. Mol Cell 12, 373–380 (2003).

51. Stojković, E. A. & Rothman-Denes, L. B. Coliphage N4 N-Acetylmuramidase Defines a New Family of Murein Hydrolases. J Mol Biol 366, 406–419 (2007).

52. Fages-Lartaud, M., Tietze, L., Elie, F., Lale, R. & Hohmann-Marriott, M. F. mCherry contains a fluorescent protein isoform that interferes with its reporter function. Front Bioeng Biotechnol 10, (2022).

53. Cámara, B. et al. T7 phage protein Gp2 inhibits the Escherichia coli RNA polymerase by antagonizing stable DNA strand separation near the transcription start site. Proc Natl Acad Sci U S A 107, 2247–2252 (2010).

54. Wah, D. A., Levchenko, I., Baker, T. A. & Sauer, R. T. Characterization of a Specificity Factor for an ATPase: Assembly of SspB Dimers with ssrA-Tagged Proteins and the ClpX Hexamer. Chem Biol 9, 1237–1245 (2002).

55. Elmore, J. R., Furches, A., Wolff, G. N., Gorday, K. & Guss, A. M. Development of a high efficiency integration system and promoter library for rapid modification of Pseudomonas putida KT2440. Metab Eng Commun 5, 1–8 (2017).

56. Snoeck, S., Guidi, C. & De Mey, M. “Metabolic burden” explained: stress symptoms and its related responses induced by (over)expression of (heterologous) proteins in Escherichia coli. Microbial Cell Factories vol. 23 Preprint at 10.1186/s12934-024-02370-9 (2024).

57. Wu, G. et al. Metabolic Burden: Cornerstones in Synthetic Biology and Metabolic Engineering Applications. Trends in Biotechnology vol. 34 652–664 Preprint at 10.1016/j.tibtech.2016.02.010 (2016).

58. Glass, D. S., Bren, A., Vaisbourd, E., Mayo, A. & Alon, U. A synthetic differentiation circuit in Escherichia coli for suppressing mutant takeover. Cell (2024) doi:10.1016/j.cell.2024.01.024.

59. Heng, Y. C., Kitano, S., Susanto, A. V., Foo, J. L. & Chang, M. W. Tunable cell differentiation via reprogrammed mating-type switching. Nature Communications 15, (2024).

60. Effendi, S. S. W. & Ng, I. S. Innovations, Challenges and Future Directions of T7RNA Polymerase in Microbial Cell Factories. ACS Synthetic Biology Preprint at 10.1021/acssynbio.5c00139 (2025).

61. Meyer, A. J., Segall-Shapiro, T. H., Glassey, E., Zhang, J. & Voigt, C. A. Escherichia coli “Marionette” strains with 12 highly optimized small-molecule sensors. Nat Chem Biol 15, 196–204 (2019).

62. Riley, L. A., Payne, I. C., Tumen-Velasquez, M. & Guss, A. M. Simple and Rapid Site-Specific Integration of Multiple Heterologous DNAs into the Escherichia coli Chromosome. J Bacteriol 205, (2023).

